# Transgene codon usage drives viral fitness and therapeutic efficacy in oncolytic adenoviruses

**DOI:** 10.1101/2020.06.19.161026

**Authors:** Estela Núñez-Manchón, Martí Farrera-Sal, Giancarlo Castellano, David Medel, Ramon Alemany, Eneko Villanueva, Cristina Fillat

**Affiliations:** Institut d’Investigacions Biomèdiques August Pi i Sunyer (IDIBAPS), Barcelona, Spain; Oncobell Program, IDIBELL, L’Hospitalet de Llobregat and VCN Biosciences, Sant Cugat del Valles, Spain; Procure Program, Institut Català d’Oncologia- Oncobell Program, IDIBELL, L’Hospitalet de Llobregat, Spain; Cambridge Centre for Proteomics, Department of Biochemistry, University of Cambridge, Cambridge, UK; Centro de Investigación Biomédica en Red de Enfermedades Raras (CIBERER), Barcelona, Spain; Facultat de Medicina i Ciències de la Salut. Universitat de Barcelona (UB), Barcelona, Spain

## Abstract

Arming oncolytic adenoviruses with therapeutic transgenes is a well-established strategy for multimodal tumour attack. However, this strategy sometimes leads to unexpected attenuated viral replication and a loss of oncolytic effects, preventing these viruses from reaching the clinic. Previous work has shown that altering codon usage in viral genes can hamper viral fitness. Here, we have analysed how transgene codon usage impacts viral replication and oncolytic activity. We observe that, although transgenes with optimised codons show high expression levels at a first round of infection, they impair viral fitness and are therefore not expressed in a sustained manner. Conversely, transgenes encoded by suboptimal codons do not compromise viral replication and are thus stably expressed over time allowing a greater oncolytic activity both *in vitro* and *in vivo*. Altogether, our work shows that fine-tuning codon usage leads to a concerted optimisation of transgene expression and viral replication paving the way for the rational design of more efficacious oncolytic therapies.

## Introduction

Cancer is the second leading cause of death globally, accounting for over 9 million deaths per year^1^. Tumours standard of care still include, in most cases, chemotherapies in combination with surgery and radiation. For many patients, these therapeutic strategies are associated with important side-effects and, unfortunately, are not always effective. An alternative treatment, able to selectively self-amplify the therapeutic effect in the tumour while non affecting healthy tissues, represents an attractive clinical approach.

Lytic viruses replicate and expand killing the infected cells. These properties of self-amplification and cell-lysis make them, theoretically, ideal for antitumour therapies. Nevertheless, for many viruses, its replication has not evolved to target tumours, so their selectivity and efficacy needs to be rationally engineered. Oncolytic virotherapy is a therapeutic approach consisting in the use of genetically modified lytic viruses, engineered to replicate in cancer cells in order to selectively kill them. Adenoviruses constitute an attractive option as oncolytic viruses, as their replication cycle is well known, present potent lytic activity, can be produced at high titres and are easily genetically modified. In the past 20 years, multiple preclinical and clinical trials have used these viruses as therapeutic agents in the treatment of different tumours^2–7^. Despite the encouraging results in terms of safety, the efficacy of adenoviral treatments still needs to be improved in order to increase the clinical usefulness of this strategy^8^.

Arming oncolytic adenoviruses (OAds) with therapeutic transgenes is an attractive approach to increase their efficacy. OAds are a highly versatile therapeutic platform that allow oncoselective expression of a wide range of therapeutic molecules such as cytokines^9^, extracellular matrix modulators^7,10^, immune checkpoint blockades (IBCs)^11^, bi-specific T-cell engagers (BiTEs)^12^ or prodrug converting enzymes^13^, allowing a multimodal tumour attack, from direct cell lysis to tumour microenvironment modulation. Moreover, expressing therapeutic molecules through OAds provides the unique opportunity to attain targeting tumours using therapeutic agents with unacceptable systemic toxicity profiles^14,15^.

A standard strategy to maximise protein production in gene therapy is the adaptation of the transgene codon usage to match the one of the host cell. Although, this could be an intuitive strategy to maximise expression of therapeutic genes in OAds, previous results have shown that adenoviral replication is tightly regulated through a balanced codon usage. In this way, viral genes avoid intergenic competition to adequately exploit translational resources for efficient replication^16^. This fact appears to be particularly important at a late phase of infection, when viral genes become highly expressed using the same finite pool of available cellular resources^17^.

In this work, we have assessed how this codon balance can be affected by transgene expression, thus impacting OAds therapeutic efficacy. This has been prompted by the observation that the codon usage of late expressed therapeutic genes in OAds currently in the clinic is in fact suboptimal when compared to host codon usage. The systematic study of the codon usage of therapeutic transgenes presented in this work, shows how transgene codon usage has not only significant *cis* effects, but also *trans* effects on other viral proteins. We provide evidence on the importance of studying armed OAds as a holistic system, rather than considering the virus and its transgenes as separate elements. We highlight the importance of developing codon usage models to reduce transgene-viral intergenic competition, thus not only maximising transgene expression, but also ensuring adequate OAds replication. In this way, concerted optimisation of transgene expression and viral replication through balanced codon usage results in more efficacious viral therapies.

## Results

### Transgene codon usage impacts oncolytic adenoviral replication

Adenoviral genes display a fine-tuned codon usage, in which highly abundant late structural proteins present codons frequently used in the human host (optimal codons) in comparison to early regulatory proteins. However, not all structural proteins have an optimal codon usage. Unlike the rest, the adenoviral fiber displays a suboptimal codon usage, which is necessary for codon balance and efficient viral replication^16^. Notably, analysis of the codon usage of transgenes expressed in OAds currently in clinical trials shows that most transgenes also use suboptimal codons, as compared to the overall codon usage in human genes (i.e., they have a codon adaptation index (CAI)^18^, a measure of directional synonymous codon usage bias, lower than the average human gene, Supplementary Fig. 1a). This would contradict the general paradigm that recommends using optimal host codons in transgenes to maximise gene expression, and suggest that this strategy could have a negative impact for therapeutic OAd fitness.

**Fig. 1:**
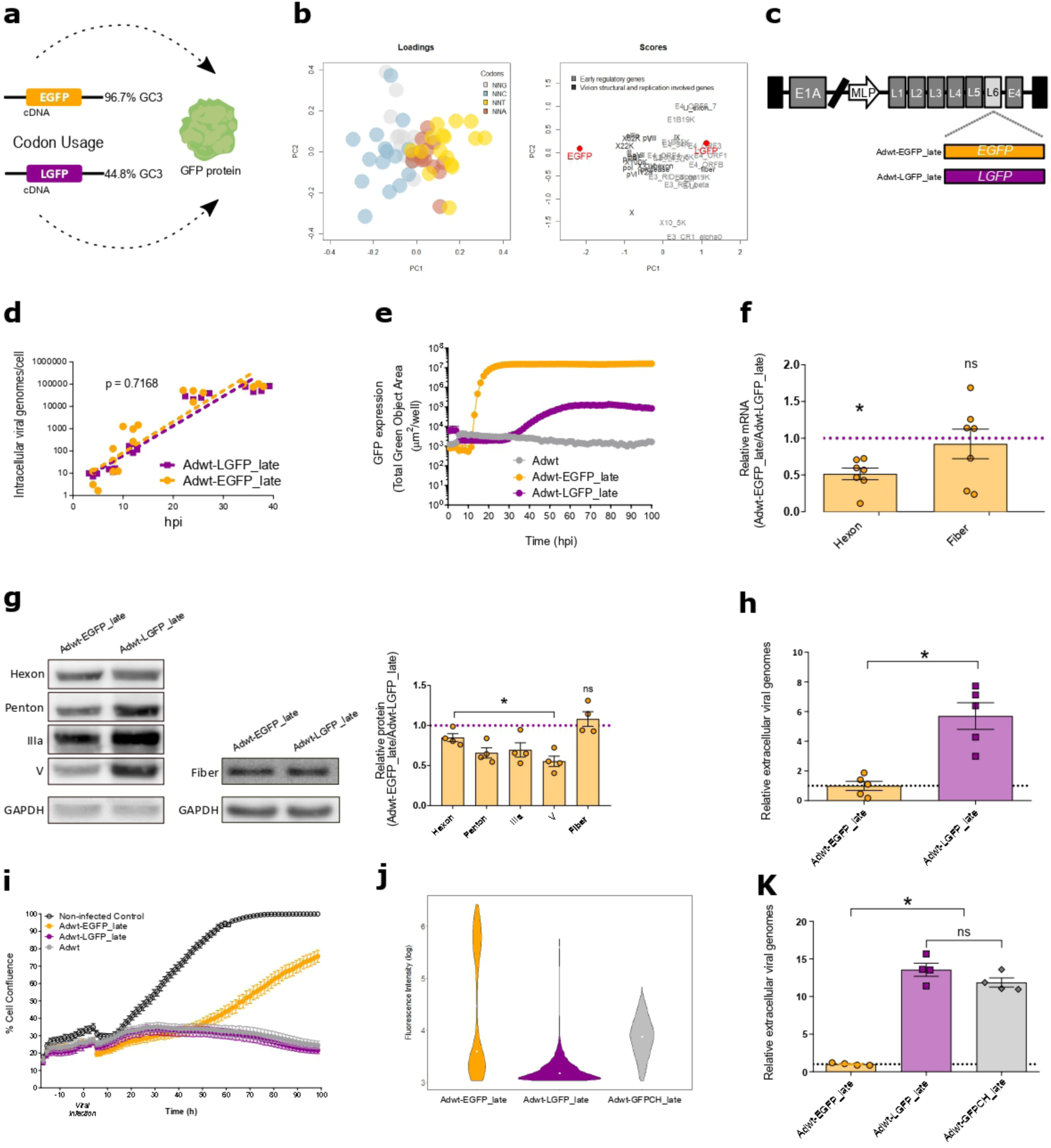
Viral fitness is affected by late transgenes in a codon-dependent manner. **a** Schematic model of GFP transgenes: the enhanced green fluorescent protein EGFP (96.7% of GC3 content), and the low codon optimised GFP LGFP (44.8% of GC3 content). **b** Transgenes codon usage evaluation by Principal Component Analysis (PCA): Loadings in the left panel showing codons coloured according to the 3^rd^ nucleotide composition; scores in the right panel showing the distribution of all viral genes as well as *EGFP* and *LGFP* transgenes in the first two principal components (PC1 and PC2). Early adenovirus regulatory genes are represented in light grey, with an increased usage of AT3 codons (orange and red spheres). Late structural and replication genes are represented in dark grey, with an increased usage of GC3 codons (grey and blue spheres). **c** Schematic representation of the whole adenoviral genome armed with *EGFP* or *LGFP* transgenes under the MLP control. **d** Intracellular viral genomes at 4, 8, 12 and 36 hpi in A549 cells infected with 10 IFU of Adwt-EGFP_late or Adwt-LGFP_late. Each dot represents an independent experimental replicates. **e** Fluorescence expression time course assay in A549 cells infected with Adwt-EGFP_late or Adwt-LGFP_late at 5 IFU and analyzed by Incucyte live cell motorization during a period of 100h. Data is represented as the mean ±SEM of five independent experimental replicates. **f** Relative viral hexon and fiber mRNA content analyzed by RT-qPCR at 36 hpi. Data is represented as the mean ±SEM; each dot corresponds to an independent experimental replicate. **g** Adenoviral late structural protein expression analysis by WB at 36 hpi. Left and central panels show representative WB. The right panel shows WB quantification. Data is represented as the mean ±SEM; each dot corresponds to an independent experimental replicate. **h** Quantification of the relative extracellular viral particles released to the supernatants of A549 cells infected with 10 IFU of Adwt-EGFP_late or Adwt-LGFP_late at 36 hpi. Data is represented as the mean ±SEM; each dot corresponds to an independent experimental replicate. **i** Proliferation assay of A549 cells infected with 5 IFU of Adwt-EGFP_late, Adwt-LGFP_late or Adwt, analyzed by Incucyte live cell motorization during a period of 100h. Data is represented as the mean ±SEM of four experimental replicates. **j** Flourescence analysis by flow cytometry 48 hpi of A549 cells infected with 0.5 IFU of Adwt-EGFP_late, Adwt-LGFP_late or Adwt-GFPCH_late. **k** Relative extracellular viral particles release analysed by qPCR at 48 hpi. Data is represented as the mean ±SEM; each dot corresponds to an independent experimental replicate. *p<0.05 (two tailed Mann-Whitney test). n.s = non-significant differences

To investigate the impact of the transgene codon usage on the oncolytic viral fitness, we selected two reporter genes: the enhanced green fluorescent protein (*EGFP)*, a broadly used GFP codon optimised to maximise its translation in human cells, and the low codon optimised GFP (*LGFP*), which mimics the CAI of the transgenes expressed by the OAds in clinical trials (Supplementary Fig. 1b). Both transgenes encode the same GFP amino acid sequence, but with different codon usage (Fig. 1a, Supplementary Data 1). *EGFP* uses mainly codons with G or C at the third base position (human optimal codons) while *LGFP* uses more frequently codons with A or T at the third base position (human non-optimal codons). The Principal Component Analysis (PCA) of the relative codon usage of both *EGFP* and *LGFP* in relation to the adenoviral genes, shows that *EGFP* gene clusters in the vicinity of the late structural and replication related genes in the first Principal Component (PC1), while *LGFP* gene clusters with the early regulatory viral genes. (Fig. 1b). PC1 discriminates viral genes according to their relative use of codons with G or C at the third base position (GC3). Positive PC1 values correlate with genes with low GC3 content (and high AT3), while negative PC1 values correlate with genes with high GC3 content (and low AT3). It has been recently reported that GC3 codons are associated with human mRNA stability and with higher translation efficiency, thereby increasing protein production^19^.

To investigate the impact of codon usage in transgene expression, *EGFP* and *LGFP* were armed as late genes under the control of the Major Late Promoter (MLP) in the wild type adenovirus 5 (Adwt) genome, generating Adwt-EGFP_late and Adwt-LGFP_late, respectively (Fig. 1c). No differences were found in the intracellular viral DNA replication at first round of infection when infecting cells at equal viral dosage (Fig. 1d), while live-cell fluorescence imaging of Awt-EGFP_late and Adwt_LGFP_late revealed that *EGFP* transgene expression was two orders of magnitude higher in relation to *LGFP*, as expected considering its higher GC3 content (i.e. its optimal codon usage) (Fig. 1e). Next, we assessed whether the transgene codon usage had any effect on viral gene expression. Analysis of the mRNA levels of the Late-phase structural hexon gene, a gene with a high percentage of GC3 codons, showed a 50% decrease in the mRNA content in cells infected with Adwt-EGFP_late in comparison with the ones infected with Adwt-LGFP_late. Interestingly, no differences in mRNA levels were found in the structural fiber gene, the only late structural gene with low GC3 content (Fig. 1f). Western Blot analysis of Late-phase hexon, penton, IIIa, V and fiber proteins further showed that Adwt-EGFP_late infected cultures displayed lower expression of structural proteins with high GC3 content, while fiber protein levels were similar between viruses (Fig. 1g). As a result of the impaired expression of many Late-phase proteins, Adwt-EGFP_late infected cultures showed six times less viral release than the Adwt-LGFP late infected ones (Fig. 1h). This difference in viral replication resulted in the impairment of Adwt-EGFP_late virus to control A549 cell proliferation (Fig. 1i). The reported differences with Adwt-EGFP_late and Adwt-LGFP_late were not observed in parallel studies conducted with transgenes inserted as early genes Adwt-EGFP_early and Adwt-LGFP_early (Supplementary Fig. 2). These results are in line with the concept of an intergenic competition for cellular resources in transgenes engineered to be expressed in the late phase of viral infection but not during the early phase, when the host resources are not yet monopolised by the virus. Altogether, these observations point out that, at the late phase of infection, transgene codon usage would not only have *cis* effects on its own expression, but also t*rans* effects on other viral proteins expressed.

**Fig. 2:**
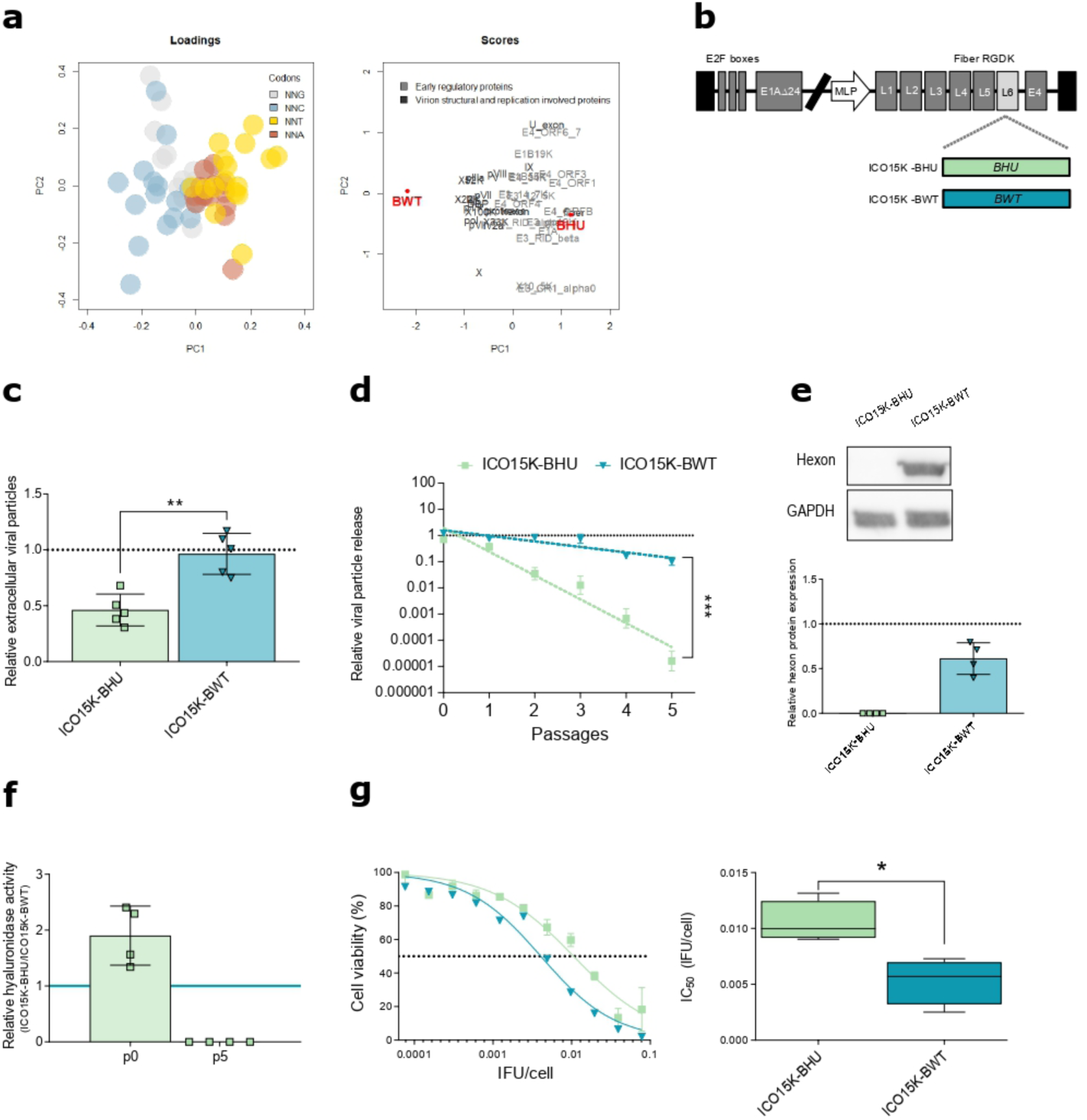
Transgene optimisation to human codon usage negatively affects viral replication and oncolysis. **a** Principal Component Analysis of the relative codon usage per amino acid of bee hyaluronidase transgenes: the left panel shows the loadings, codons characterised according to the 3^rd^ nucleotide composition, the right panel shows the hyaluronidase transgenes (red) in relation to adenoviral genes. **b** Schematic representation of ICO15K-BHU and ICO15K-BWT virus, armed with codon humanised bee hyaluronidase (*BHU*) and codon wild type bee hyaluronidase (*BWT*), respectively. **c** qPCR relative quantitation of the extracellular viral particles released to the supernantants of PANC-1 cells infected with 5 IFU of VCN-01, ICO15K-BHU or ICO15K-BWT at 72 hpi,. The dashed line represents VCN-01 values. Data is represented as the mean ±SEM; each dot corresponds to an independent experimental replicate. **p<0.01 (two tailed Mann-Whitney test). **d** qPCR relative quantification of viral release in the supernatants of PANC-1 cells infected with ICO15K-BHU, ICO15K-BWT or ICO15K-BAd at several rounds of infection. The dashed line represents VCN-01 values. Data is represented as mean ± SEM of four independent experiments. Differences between slopes were analysed using F-test for nonlinear models. *** p<0.001. **e** Hexon protein analysis by Western Blot at passage 5. Upper panel: representative WB. Lower panel: WB quantification of 4 independent replicates. The dashed line represents VCN-01 values. **f** Turbidimetric quantification of hyaluronidase activity in the supernatants of PANC-1 cells infected with ICO15K-BHU and ICO15K-BWT at passages p0 and p5. **g** *In vitro* oncolytic activity assay in PANC-1 cells. Cells were infected with a dose range of ICO15K-BHU and ICO15K-BWT and their viability was measured 7 days PI by MTT assay. Viability curves are represented in the left panel as mean ±SEM of at least four independent experiments. The right panel represents IC_50_ values as box plot of four independent experiments. *p<0.05 (two tailed Mann-Whitney test).

For an effective therapeutic output, it is necessary to properly balance the viral lytic activity and therapeutic transgene expression. Therefore, we next evaluated whether tuning transgene codon usage could allow a better transgene expression, without compromising viral replication. We designed three new chimeric transgenes encoding GFP (*CH1, CH2* and *CH3*) by combining *EGFP* and *LGFP* sequences (Supplementary Fig. 3a-b) to achieve intermediate codon usage optimization (Supplementary Fig. 3c-f). GFP expression analysis upon transfection evidenced that the levels of fluorescence of the different transgenes were highly dependent on their GC3 content (Supplementary Fig. 1f-d). *CH1* transgene was selected and armed under the MLP, generating Adwt-GFPCH_late. Importantly, Adwt-GFPCH_late infected cultures presented higher transgene expression capacity than cell cultures infected with Adwt-LGFP_late (Fig. 1j) without impairing viral fitness (Fig. 1k). In this way, our results suggest that oncolytic activity and late transgene expression can be optimised through the balancing of codon usage.

**Fig. 3:**
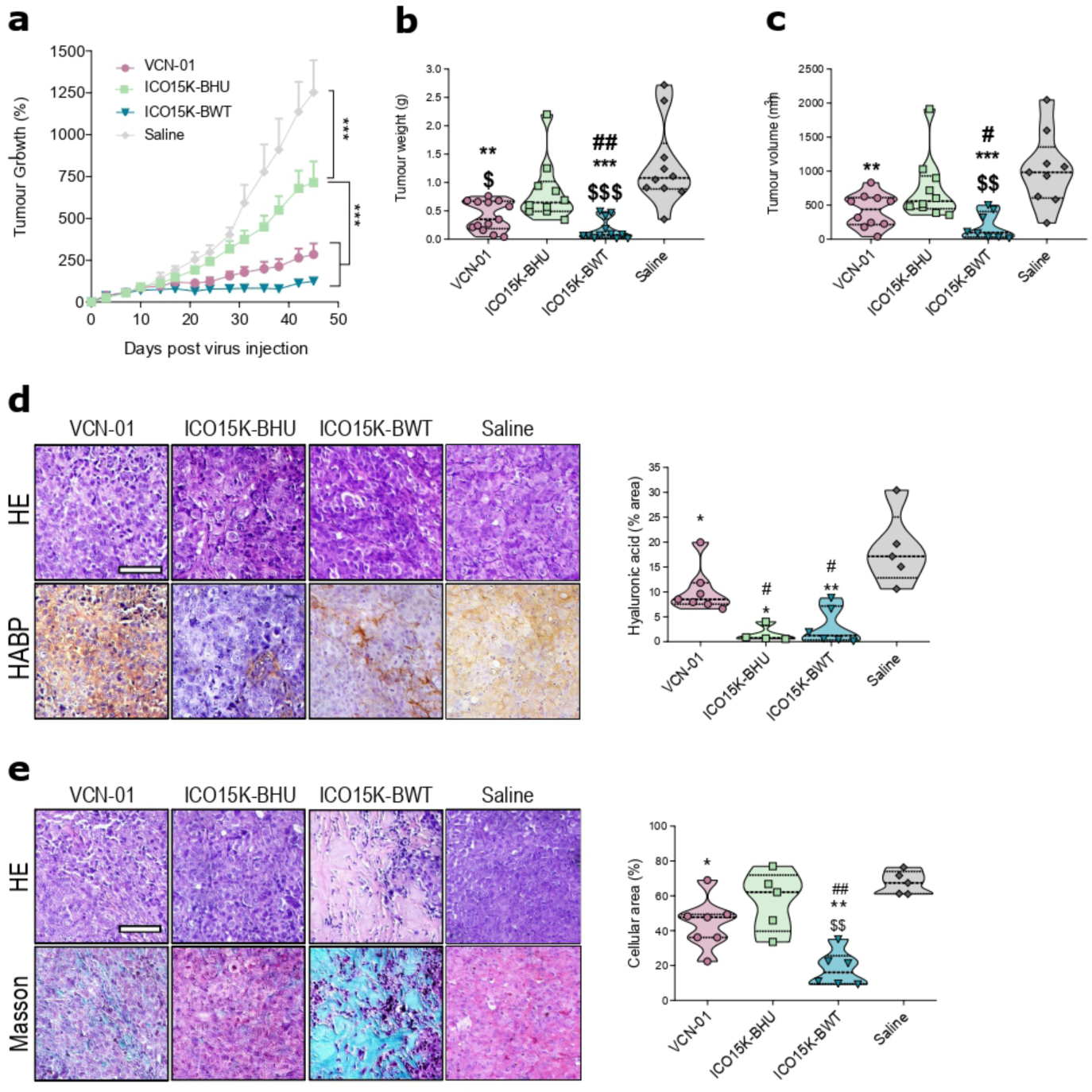
Balanced virus and transgene expression leads to optimal therapeutic index response. **a** *In vivo* tumour growth assay in mice bearing subcutaneous PANC-1 tumours. Animals were intravenously treated with saline solution or 4··10^10^ vp/animal of either VCN-01, ICO15K-BHU or ICO15K-BWT (n≥6 animals/group). Follow-up of tumour volumes is represented as mean of percentage of growth ±SEM. **b** Tumour weight at end-point. **c** Tumour volume at end-point. **d** HA acid staining quantitation of deparaffinised tumour sections with HABP. Scale bar 100 μ m. Left panels: representative images of HA staining. Right panels: HA acid quantification representing stained area in at least 5 fields from the tumour sections of at least 3 different mice. **e** Tumour cellularity measured by Masson staining. Scale bar 100 μ m. Left panels: representative images of HA staining. Right panels: HA acid quantification. *p<0.05, **p<0.01, ***p<0.001, # p<0.05, ## p<0.01, $ p<0.05, $$ p<0.01, $$$ p<0.001 * represents statistical differences in relation to saline group; # represents statistical differences in relation to saline VCN-01 group, $ represents statistical differences in relation to ICO15K-BHU group.

### Sustained therapeutic transgene expression depends on viral fitness

To assess the importance of balancing the lytic activity of the virus and the transgene expression in a therapeutic context, we evaluated the impact of transgene codon usage in the design of oncolytic viruses. We selected two versions of the bee (*Apis mellifera*) hyaluronidase, with different codon usages, as therapeutic transgenes (Supplementary Data 2). Previous oncolytic virus armed with the human hyaluronidase (*PH20* gene expressed by the VCN-01 virus) showed that the expression of this enzyme allows tumour microenvironment remodeling, making tumour cells more accessible to anti-tumour treatments^7^. Since bee hyaluronidase presents higher enzymatic activity than its human orthologue (Supplementary Fig. 4), we reasoned that arming oncolytic viruses with bee hyaluronidase could enhance their antitumour activity. We selected a humanised version of the bee hyaluronidase (*BHU*), modified following standard codon humanization algorithms (performed by Genscript) (Supplementary Fig. 5) displaying high GC3 content (Fig. 2a), and the wild type bee hyaluronidase (*BWT*) with a suboptimal codon usage for its expression in human cells (Supplementary Fig. 5), and a GC3 content similar to that of the *CH1* gene (Fig. 2a). We used the viral platform ICO15K (the same one of the VCN-01 virus, currently in clinical trials) to express the *Apis mellifera* hyaluronidase transgenes with different codon usages (Fig. 2b). ICO15K is a E1-Δ24 engineered adenovirus with four E2F and one sp-1 binding sites in the E1A promoter and the RGDK motif replacing the KKTK glycosaminoglycan binding domain in the fiber shaft^7,20^.

**Fig. 4:**
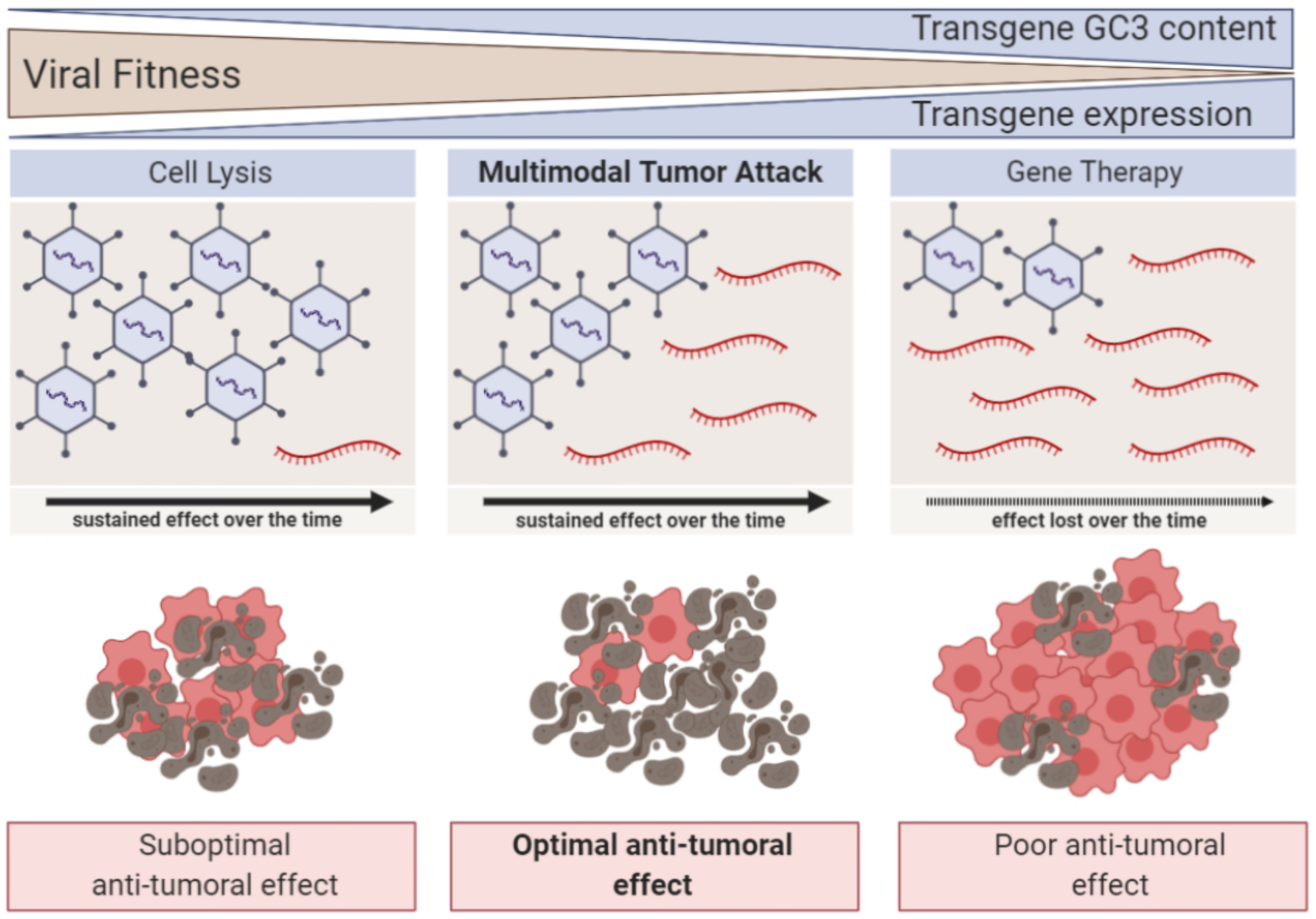
Transgene codon usage drives therapeutic efficacy of armed oncolytic adenoviruses. Graphical representation of transgene codon usage impact on viral fitness and oncolytic virotherapy therapeutic efficacy.

As expected, the virus expressing the human optimised bee hyaluronidase, ICO15K-BHU, showed a significant impairment of viral fitness, similar to the one of the Adwt-EGFP_late virus (Fig. 2c and Supplementary Fig. 6a). Viral replication impairment was further evidenced after several rounds of infection (Fig 2d). Indeed, after 5 rounds of consecutive infections, the ICO15K-BHU virus became extinct (Fig. 2d-e). Consequently, *BHU* transgene hyaluronidase activity was not detected at passage 5, despite displaying the highest enzymatic activity at passage 0 (Fig. 2f). Cytotoxicity assays carried out in pancreatic ductal adenocarcinoma (PDAC) cell models PANC-1 (Fig. 2g), MIA PaCa-2 and NP18 (Supplementary Fig. 6b-c) evidenced a significant decrease in the oncolytic capacity of ICO15K-BHU compared to ICO15K-BWT, in concordance with its impaired viral replication. Collectively, our results show that viral fitness of oncolytic adenoviruses armed with canonically humanized transgenes negatively impact viral replication. However, the therapeutic activity of the virus can be rescued through a codon optimization that takes into account the effect of the transgene on virus replication.

### Preserving viral lytic activity is key to maximise antitumoural efficacy

To investigate how the interplay between transgene codon optimization and viral lytic capacity impacts therapeutic efficacy, we treated athymic mice bearing subcutaneous PANC-1 and MIA PaCa-2 tumours with a collection of OAds armed with different hyaluronidases. Both pancreatic cell lines generate tumours with high density of stroma, mimicking pancreatic tumours. In these conditions, viral spread is impaired and the antitumoural activity of OAds is limited. Therefore, arming OAds with potent hyaluronidase transgenes should boost the therapeutic activity of these viruses. However, we observed that the virus expressing the human optimised bee hyaluronidase, ICO15K-BHU, was the least efficient reducing tumour growth, size and weight (Fig. 3a-c and Supplementary Fig. 7 a-c). Thus, despite displaying the highest hyaluronidase activity, which led to a notable decrease of hyaluronic acid (Fig. 3d and Supplementary Fig. 7d), tumours treated with this virus maintained high cellularity (Fig. 3e and Supplementary Fig. 7e), demonstrating that, prioritising high transgene activity is insufficient for controlling tumour progression if the viral lytic activity is impaired. This is consistent with the results from our previous *in vitro* analyses (Fig. 2). On the contrary, treatment with the virus expressing the bee hyaluronidase gene with a sub-optimal codon usage, ICO15K-BWT, resulted in the highest antitumour activity (Fig. 3a-c and Supplementary Fig. 7a-c). When comparing the therapeutic activity of ICO15K-BWT with the VCN-01 virus, which expresses the human hyaluronidase and is currently in clinical trials, the virus with the bee hyaluronidase showed increased antitumour activity, both by efficiently remodelling the tumoural microenvironment (Figure 3d-e) and by controlling tumoural growth (Fig. 3a-c). In tumours derived from a cell line with reduced viral sensitivity (ie. MIA PaCa-2 and Supplementary Fig. 6b), ICO15K-BWT treatment controlled tumour growth (similarly to VCN-01) (Supplementary Fig. 7a-c) and enhanced tumoural microenvironment remodelling (Supplementary Fig. 7d), although it was not capable of reducing cellularity in the remaining tumour mass (Supplementary Fig. 7e). The fact that all viruses expressing *Apis Mellifera* hyaluronidase showed a significant decrease of hyaluronic acid, when compared to the VCN-01 virus expressing the human hyaluronidase (Fig. 3d and Supplementary Fig. 7d) is in line with our observations that bee hyaluronidase has a superior enzymatic activity (Supplementary Fig. 3). Altogether our results reveal that preserving viral lytic activity is key to maximise the therapeutic output of the virus and highlight the importance of adequately balance viral fitness and transgene expression to optimise treatment efficacy.

## Discussion

Oncolytic virotherapy depends on the viral capacity to successfully replicate, lyse the tumour cell and generate new viral particles by taking advantage of cellular resources. Arming oncolytic viruses with therapeutic transgenes is required to enhance the antitumour efficacy of viral therapies. Surprisingly, our analysis shows that the therapeutic genes that have been armed in OAds currently in clinical trials, have a codon usage that is not optimised to maximise their expression in human cells. Considering that codon optimisation is a commonly used strategy to maximise gene expression, we speculated that the bias towards non-optimised transgenes in armed OAds reaching the clinic was not random but respond to a biological mechanism not yet known. With the aim to understand it, we systematically studied the impact of the codon usage of therapeutic transgenes in OAd antitumour activity. Using both reporter genes and transgenes of therapeutic interest, we found that their codon usage severely impacts viral replication, and that viral replication impairment is directly correlated to the expression of the transgene. The viral-transgene competition diminishes viral replication over time, thereby extinguishing the virus in as few as five replication cycles, and consequently, abrogating therapeutic transgene expression. We found that preserving viral lytic activity over transgene expression is the key element to maximise the antitumour efficacy of the OAds.

Interestingly, we found that the viral-transgene competition is less important when expressing the transgenes during the early stages of viral replication (Supplementary Fig.2). This would be explained because the adenoviral early regulatory genes have an AT3 biased codon usage that may prevent the competition with GC3 rich optimised transgenes. Moreover, we speculate that differences in competition could be due to the ability of the cell to adapt to different translational demands in early stages of infection, while during the late phases of the viral replication cycle, the cellular machinery is barely able to satisfy viral demand. In fact, we have recently shown that during the late phase of infection, cellular translational machinery is extensively and exclusively exploited by the virus, leading to a fine-tuned balance between the supply and demand of a limited pool of cellular resources^16^. In the same line, the data presented in our work suggests that interfering with this fine-tuned translational equilibrium by expressing therapeutic transgenes could generate a translational imbalance with a dramatic effect on the therapeutic activity of OAds. Even if this could suggest that expression of transgenes in the early phases of infection could be preferable, this strategy is not always possible. Transgenes encoding toxic proteins can impact viral replication or affect viral DNA synthesis^8,21^. Moreover, if the transgene length is close to the viral encapsidation limit, the usage of exogenous promoters, required for early expression, will further increase the length of the construct, and further hinder viral assembly. Arming therapeutic transgenes under the control of the viral major late promoter can overcome or at least attenuate these effects. However, in that case, the intergenic competition of the virus and transgene observed in our work has to be considered. To facilitate so, we have created the web tool ONATRY (Oncolytic Adenovirus Transgene analYzer), that recapitulates two key analysis: transgene GC3 content analysis in human host context and codon usage PCA in relation to adenoviral genes. The tool is available at http://172.21.104.19:3838/sample-apps/ToolPCAadenovirus/. We think that the web tool we have developed could contribute to identify incompatible transgene codon-usages and thus help to improve the viral design.

Traditionally, due to the limited packaging capacity of viruses, transgene length has been considered as the main constraint when designing armed OAds^22^. Our data prompt us to reconsider this strategy to design oncolytic viruses. In particular, it highlights the need to consider the interplay between adenoviral replication and transgene expression. This would require a paradigm change in which OAds and transgenes are no longer seen as two separate elements that can be simply combined, and instead consider them as a holistic system. Thus, apparent poor transgene expression as a result of a suboptimal codon usage can paradoxically end up maximising transgene expression over time, due to a better viral replication combined with the inherent auto-amplification of the treatment (Fig. 4). Altogether, our results suggest that oncolytic virus design should abandon the traditional dogma of enhancing transgene expression as much as possible and consider the efficacy of oncolytic therapies as the combined effect of transgene expression and viral replication, where preserving viral lytic activity is key to maximise antitumour efficacy.

In more general terms, our work points to the intriguing hypothesis that many oncolytic viruses may have never reached clinical trials because the deleterious effect of the expression of a codon-optimised transgene could have prevented correct viral replication. We think that by fine tuning the codon usage of the therapeutic transgene, many of these viruses could be rescued, directly impacting the efficacy and availability of novel OAds for cancer treatment. Our findings represent a new step forward in the understanding of the interplay between transgenes and therapeutic viruses, and could guide new strategies fostering the use of oncolytic adenoviruses in the clinic.

## Methods

### Cell lines

Cell lines PANC-1 and MIA PaCa-2 (human pancreatic carcinoma), A549 (human lung carcinoma) and HEK293 (embryonic kidney) were obtained from the American Type Culture Collection (ATCC, Manasas, VA); human pancreatic adenocarcinoma cell line NP18 was established and kindly provided by Dr. Gabriel Capellà (ICO-IDIBELL, Barcelona Spain). All cell lines were maintained in Dulbecco’s modified Eagle’s medium (Gibco-BRL) supplemented with 10% fetal bovine serum, penicillin (100 μ g/ml) and streptomycin (100 μ g/ml) (Gibco-BRL), and maintained in humidified atmosphere of 5% CO2 at 37ºC.

### Expression plasmids generation and transfection

*EGFP* and *LGFP* genes were amplified using the primers with BamHI and EcoRI restriction sites (Supplementary Table 1, primer sets 1 and 2) and amplified fragments were digested with the corresponding restriction enzymes. Digested fragments were resolved in agarose gels and purified with the QIAquick Gel Extraction Kit (Qiagen) according to manufacturer’s instructions, CH1, CH2 and CH3 genes were purchased in the form of gBlock (IDT). The corresponding fragments were inserted in miRVec expression plasmid (restricted with the same enzymes) by ligation with T4 ligase (Roche) according to manufacturer’s instructions. All constructs were tested by Sanger-DNA sequencing at Beckman Coulter Genomics. Hek293T were transfected with CalPhos (Clontech) following the manufacturer’s instructions.

Similarly, *PH20* and BWT genes were amplified and flanked with Age-I and Not-I restriction sites with corresponding primers (Supplementary Table 1, primer sets 3 and 4). The amplicons were digested with Age-I and Not-I enzymes and the subsequent digested fragments were resolved by agarose electrophoresis and purified with Monarch DNA gel extraction kit (New England Biolegends) according to manufacturer’s protocol. The corresponding fragments were inserted in pGT4082 expression plasmid (restricted with the same enzymes) by ligation with T4 ligase (New England Biolegends) according to manufacturer’s instructions.

### Codon usage analysis

Codon usage frequencies were analyzed using the Sequence Manipulation Suite (). The relative codon usage was obtained normalising the codon usage of each codon by every synonymous codon for each aminoacid. The Codon Adaptation Index (CAI) of each sequence and the percentage of codons with G or C at the third base position (GC3%) were calculated using the CAIcal server from http://ppuigbo.me/programs/CAIcal/^23^. *Homo sapiens* codon usage was extracted from the Codon Usage Database available at http://www.kazusa.or.jp/codon/. The human genes analysed were selected from the Tissue-specific Gene Expression And Regulation database at the Johns Hopkins University. Available transgene sequences from the virus in clinical trials were obtained from the correspondent patents: VCN-01 (EP2428229B1), NG-641 (WO2018041838A1), NG-350A (WO2018220207A1), LoAd703 (WO2015155174A1), Ad5-yCD/mutTKSR39rep-hIL12 (WO2007087462A2), ONCOS-102 (WO2010072900), CG0070 (WO2010072900). PCA, CAI and GC3% analysis were represented using R v3.2.3 software.

### Adenovirus generation and titration

Adwt and ICO15K backbones were previously generated as described in^20,24^. GFP and hyaluronidase transgenes were introduced after fiber (L5 gene) under a IIIa splicing acceptor (IIIaSA)^25^ in Adwt and ICO15K backbones^20,24^, to generate adenoviral genomes with transgenes inserted as late units (L6). Adenoviral genomes with GFPs inserted as an early gene were generated introducing the transgenes under the control of the constitutive citomegalovirus promoter (CMV) in the region between the E4 gene and the right ITR. In all cases, transgenes were incorporated following an adapted recombineering protocol based on homologous recombination in bacteria^26,27^. Recombination fragments were obtained amplifying the transgenes with primers with the corresponding homologous sequences (Supplementary table 1, primer sets 5 to 8). *BHU* transgene was purchased as a gBlock IDT) flanked with the corresponding homologous regions. Plasmids were transfected into HEK293 cells to obtain a first round of viral particles. All viruses were propagated in A549 cells and purified using cesium chloride double-gradients following standard techniques^28^. Adenoviral titres were calculated based on optical density (using viral particles [vp]/ml) and on viral infectious units (IFU/ml) as previously described^16^.

### Viral genomes quantification

Viral DNA was obtained from the supernatants of infected cells using Norgen’s Blood DNA Isolation Mini Kit (NORGEN BIOTEK CORP.). Adenoviral DNA content was quantified by qPCR using LightCycler 480SYBER Green I Master Mix (Roche Diagnostics), (Supplementary Table 1, primer set 9). All qPCR reactions were done in a ViiA 7 Real-Time PCR System (Applied Biosystems).

### cDNA synthesis and real-time qPCR

Total and viral RNA were obtained from infected cells using RNeasy Mini Kit (Qiagen), 500 ng were reverse transcribed using PrimeScript RT-PCR Kit (Takara), and the qRT-PCR analysis was performed using LightCycler 480SYBER Green I Master Mix (Roche Diagnostics), (Supplementary Table 1, primer sets 10 and 11). qRT-PCR results were normalised to the beta-actin expression (Supplementary Table 1, primer set 12). qPCR reactions were done in a ViiA 7 Real-Time PCR System (Applied Biosystems).

### Western Blot analyses

Total protein extracts were obtained using a lysis buffer (50 mM Tris-HCl [pH 6.8], 2% SDS, and 10% glycerol) containing 1% of Complete Mini Protease Inhibitor (Roche). Cell lysates were boiled (10 min at 98ºC) and centrifuged (5 min at 16.000 g) to eliminate insoluble cellular debris. Protein concentration was determined by BCA Protein Assay kit (ThermoFisher Scientific). Fifteen μ g of protein were resolved by electrophoresis on a 10% acrylamide gel and transferred to nitrocellulose membrane by standard methods. Membranes were incubated with Anti-Adenovirus Type 5 Capsid antibody (1:1000; Abcam) for hexon, penton, IIIa and V proteins detection and with Adenovirus Fiber [4D2] antibody (1:200; GeneTex) for fiber detection. Protein labeling was detected using HRP-conjugated antibodies and visualized in the image reader LAS4000 (Fujifilm). All protein expression data was normalized to GAPDH protein expression.

For hyaluronidase quantification, supernatants from HEK293 cells were collected 5 days after transfection with pGT4082-hyaluronidase expressing plasmids. Equivolumes of respective transfections (30DL) were resolved in a 12% acrylamide gel together with a standard curve of commercial purified recombinant His-tagged hPH20 (Acro Biosystems, PH0-H5225). The gel was transferred to nitrocellulose membrane and overnight incubation with Anti-HisTag antibody (1:4000, Dianova) was performed. Then a secondary antibody anti-mouse-IgG HRP (1/2000, Dako) was used to reveal and subsequent visualisation of the western blot in the Chemi-doc (BioRad).

### Cytotoxicity

Cells were seeded in triplicate and infected with serial dilutions of each virus. At 4 h post infection, virus-containing medium was replaced with fresh medium. Cell viability was measured 7 days post infection by a colorimetric assay following the manufacturer’s instructions (MTT Ultrapure; USB).

### Flow cytometry

Evaluation of fluorescence intensity by flow cytometry was performed using Attune Acoustic Focusing Cytometer (Applied Biosystems) and analyzed using FlowJo 8.7 for Macintosh.

### Hystological analysis

PANC-1 and MIA PaCa-2 tumours were fixed in a 4% paraformaldehyde solution overnight, and embedded in paraffin. Standard Masson and hematolycin and eosin staining were performed in 5 μm sections. For hyaluronic acid staining, permeabilization and blockade was done using PBS with 0.3% triton, 10% FBS and 1% BSA. HABP (Sigma) at 5μ g/ml was incubated overnight and the HA labeling was detected using Vectastain® ABC kit (Vector Laboratories) and DAB (Vector Laboratories). Hematoxylin counterstaining was performed for 2 min at room temperature (RT).

### Hyaluronidase activity detection - Turbidimetric assay

Supernatants from infected or transfected cells were mixed with a HA (Sigma, St Louis, MO) solution in phosphate buffer (pH 6.0) and incubated for 14h at 37 °C. Standards with known concentrations of hyaluronidase were incubated in parallel to generate standard curves. Five volumes of acid albumin (24 mM sodium acetate, 79 mM acetic acid, 0.1% bovine albumin [pH 3.75]) were added to all samples and incubated for 10 min at RT. The hyaluronidase activity was measured by light absorbance at 600nm. Absorbance values of supernatants were used to calculate hyaluronidase activity by extrapolation to the standard curves.

### Antitumoural *in vivo* study

Subcutaneous tumours were generated in 6-7-week-old male Athymic Nude Foxn1nu/nu mice (ENVIGO) by injecting 2×10^6^ MIA PaCa-2 or PANC-1 cells embedded in Matrigel 1:1 (BD Biosciences) into each flank. Tumours were measured at least twice weekly, and the tumoural volumes calculated using the formula V = D × d^2^ × π ÷ 6 V. Mice were randomly assigned to either group for treatment. Viruses (4·10^10^ vp/animal) were administered intravenously in physiological saline solution once tumours achieved a median volume of 100 mm^3^. The experiment remained blinded until the conclusion of the study. Animals were euthanized 45-48 days after virus administration, and tumours were collected and flash-freezed in liquid nitrogen. All animal procedures met the guidelines of European Community Directive 86/609/EEC and were approved by the ethical committee (CEEA-University of Barcelona) and by the local authorities of the Generalitat de Catalunya.

### Statistical Analysis

Statistical analysis was performed on GraphPad Prism v8.0.1 (GraphPad Software). Unless specified, statistical differences were evaluated using a 2-tailed non-parametric Mann-Whitney test. The level of significance was set as p < 0.05.

The *in vivo* tumour growth statistical analysis was evaluated using a linear mixed-effect model with the lme4 package in R v3.2.3. We associated a random-effects term with the day of measurement^29^. Statistical differences were evaluated using a multiple comparison of means by Tukey’s contrasts.

### Web Tool

ONATRY is a graphical user interface for analysing adenoviral transgene codon usage suitability. ONATRY represents transgenes in the context of viral genes according to their GC content and codon usage via Principal Component Analysis (PCA). It is implemented in R (version 3.6.3) and encapsulated in a Shiny application (https://shiny.rstudio.com/). Users can access the tool via a web interface (http://172.21.104.19:3838/sample-apps/ToolPCAadenovirus/).

Our Shiny application pipeline provides two different ways to input data for the analysis. First, a file in fasta format can be uploaded (through a ‘fileInput’ button). Second, there is the possibility to paste sequences in fasta format (via ‘textAreaInput’). ONATRY provides two visualization tools: a density plot and a PCA representation for exploratory analysis. The current implementation depends on the following R packages: shiny, coRdon, dplyr, rgl, and progress.

## Acknowledgements

This work was supported by grants to CF and RA from the Spanish Ministry of *Economia y Competitividad* BIO2017-89754-C2-1R and 2R co-financed by Fondo Europeo de Desarrollo Regional (FEDER), with partial support from the *Generalitat de Catalunya* SGR17/861 and 2017SGR449. CIBER de *Enfermedades Raras* is an initiative of the ISCIII. We also acknowledge the support of the Spanish Adenovirus Network (AdenoNet, BIO2015-68990-REDT) and the CERCA Programme/Generalitat de Catalunya. This work was developed at the Centro Esther Koplowitz, Barcelona, Spain. We are indebted to the Banc de Tumours-Biobank core facility of the Hospital-Clínic-IDIBAPS for technical help (work supported by the Xarxa de Bancs de Tumours de Catalunya - XBTC). We would like to thank Dr Maria Marti-Solano for her feedback during the manuscript writing process.

## Author Contributions

EN-M, EV and CF conceived and designed the experiments. EN-M conducted most of the *in vitro* and *in vivo* experiments. MF-S contributed with the hyaluronidase assay and with the experiments of hyaluronidase expressing plasmids. RA contributed with the generation of ICO15K-BHU and experimental suggestions. GC and DM developed the web tool. EN-M, EV and CF wrote the manuscript, with input from all other authors.

