## Supplemental Information for "Transgene codon usage drives viral fitness and therapeutic efficacy in oncolytic adenoviruses"

**Supplementary Information**

**a**

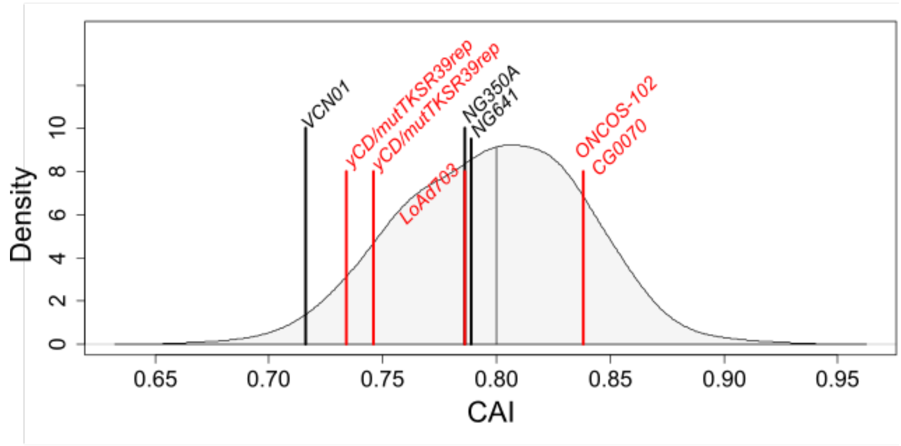


**b**

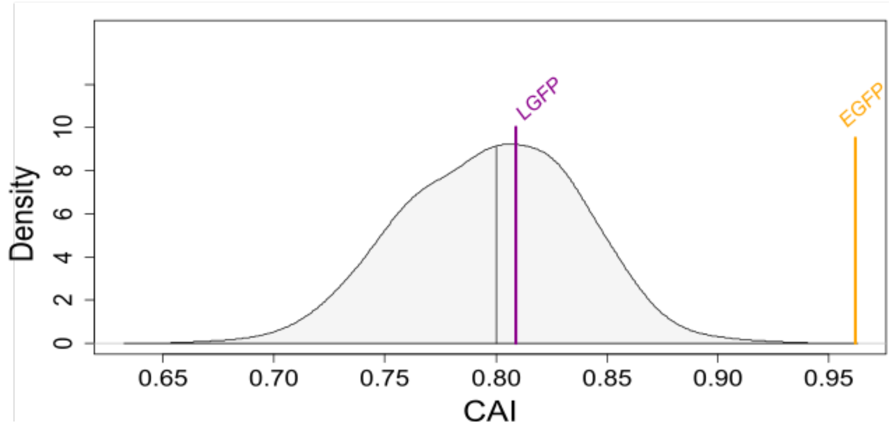


**Supplementary Fig. 1: Transgenes expressed in OAds currently in clinical trials use suboptimal codons.**

**a** Codon Adaptation (CAI) analysis of clinical trial OAds transgenes in comparison with 700 randomly selected human proteins (grey) with a men CAI value of 0.80. Red and black vertical lines correspond to the CAI value of clinical trials OAds transgenes cassettes (with available sequence) expressed in early and in late viral phase, respectively.

**b** CAI analysis of the enhanced green fluorescent protein (*EGFP)* and the low codon optimised GFP (*LGFP*).

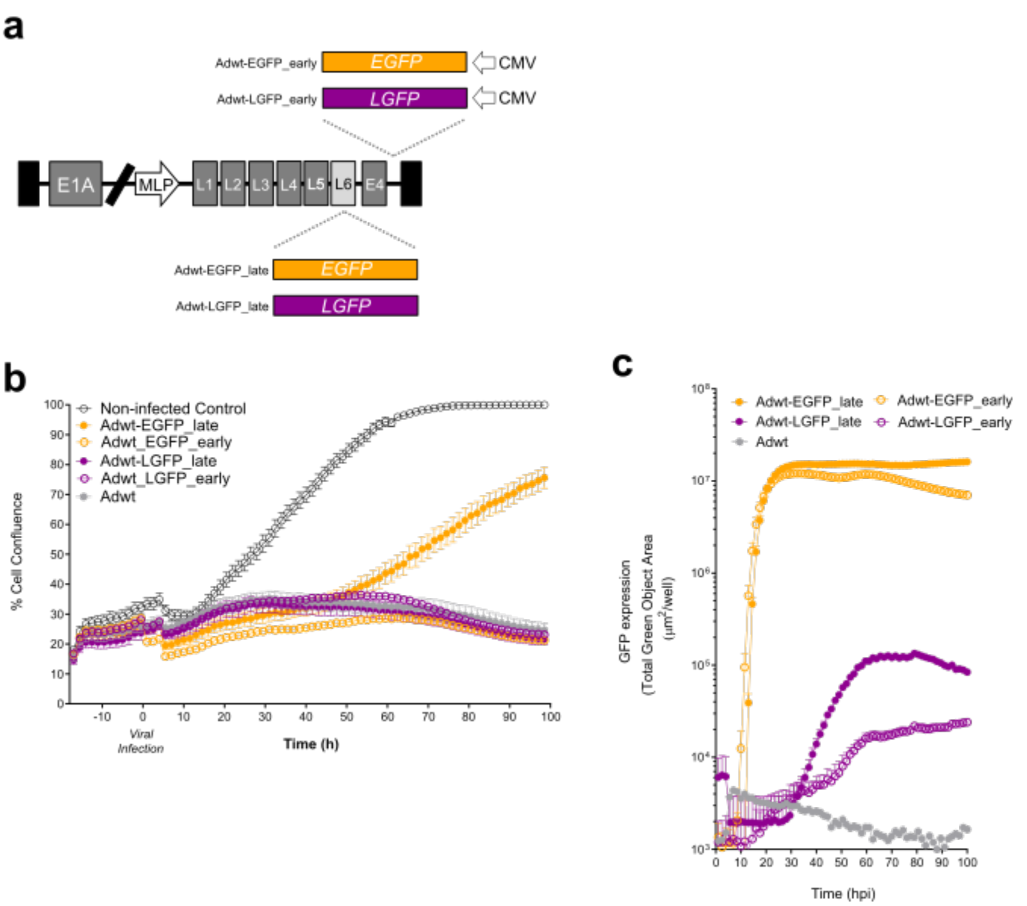


**Supplementary Fig. 2: Transgene codon usage does not impact viral fitness when expressed at early phase of infection.**

**a** Schematic representation of the whole adenoviral genome armed with EGFP or LGFP transgenes in late (under the MLP control downstream the L5-Fiber transcription unit) and in early (under the control of CMV constitutive promoter inserted between E4 and the right ITR).

**b** Proliferation assay in A549 cells infected with 5 IFU of Adwt-EGFP_late, Adwt-LGFP_late, Adwt-EGFP_early, Adwt-LGFP_early or Adwt, analyzed by Incucyte live cell motorization during a period of 100h. Data is represented as the mean ±SEM of four experimental replicates.

**c** Fluorescence assay in A549 cells infected with 5 IFU of Adwt-EGFP_late, Adwt-LGFP_late, Adwt-EGFP_early or Adwt-LGFP_early, analyzed by Incucyte live cell motorization during a period of 100h. Data is represented as the mean ±SEM of five independent experimental replicates.

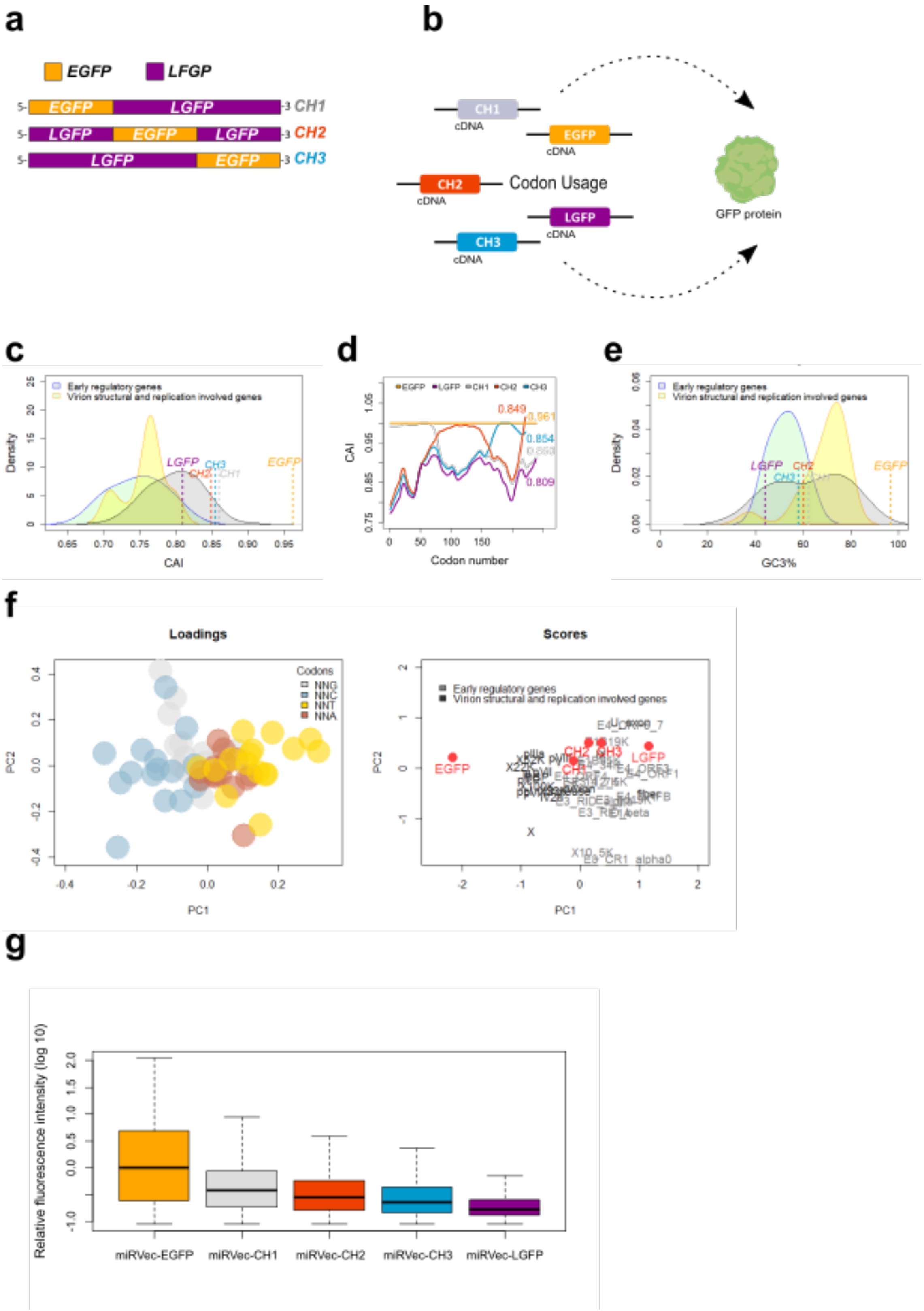


**Supplementary Fig. 3: Hybrid transgenes obtained from EGFP and LGFP sequences express GFP proportionally to their codon optimization in terms of GC3 content.**

**a** Schematic model of chimeric GFP transgenes *CH1*, *CH2* and *CH3* design.

**b** Schematic model of all GFP transgenes: the enhanced green fluorescent protein EGFP, the low codon optimised green fluorescent protein LGFP and the chimeric GFPs CH1, CH2 and CH3.

**c** Global Codon Adaptation Index (CAI) analysis according to the human codon usage of GFP transgenes in comparison to early regulatory adenoviral genes (in blue) and to virus structural and replication involved genes (in yellow).

**d** CAI along the sequence; line pattern represents smoothened CAI values along the sequence and the numeric value corresponds to the global CAI value for each sequence.

**e** Codon usage optimization analysis in terms of GC3% of GFP transgenes in comparison to early regulatory adenoviral genes (in blue) and to virus structural and replication involved genes ( in yellow).

**f** Transgenes codon usage evaluation by Principal Component Analysis (PCA): Loadings in the left panel showing codons coloured according to the $3^{rd}$ nucleotide composition; scores in the right panel showing the distribution of all viral genes as well as *EGFP, LGFP, CH1, CH2* and *CH3* transgenes in the first two principal components (PC1 and PC2). Early adenovirus regulatory genes are represented in light grey, with an increased usage of AT3 codons (orange and red spheres). Late structural and replication genes are represented in dark grey, with an increased usage of GC3 codons (grey and blue spheres).

**g** Fluorescence analysis by flow cytometry of 293T cells 48h post-transfection with miRVec-EGFP, miRVec-CH1, miRVec-CH2, miRVec-CH3 and miRVec-LGFP expression plasmids. Data is represented as box plot of three independent experiments.

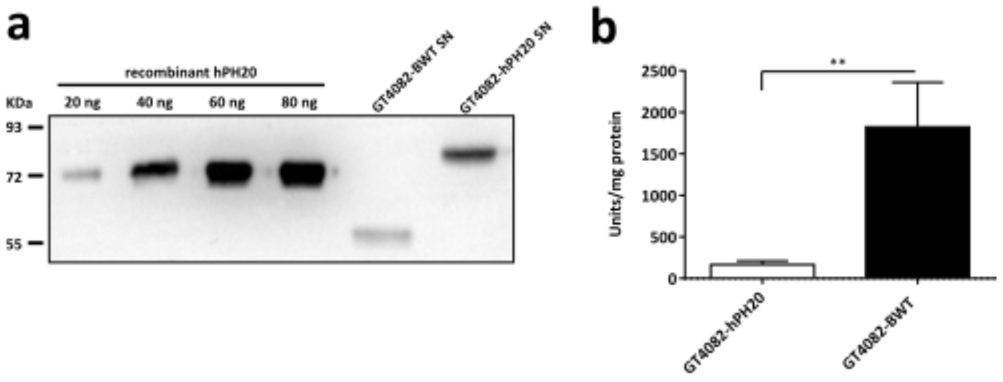


**Supplementary Fig. 4: Bee hyaluronidase (BWT) activity is higher than human hyaluronidase PH20.**

**a** Equivolumes of supernatants from 5-days HEK293 cells transfected with GT4082-BWT or GT4082-hPH20 plasmids were assessed by anti-HisTag Western Blot. A commercial purified recombinant His-tagged hPH20 (Acro Biosystems, PH0-H5225) was used as a positive control to perform a standard curve ranging from 20 to 80 ng in order to quantify the samples.

**b** Hyaluronidase activity in the supernatants of the 5 days HEK293 transfected cells was analysed by turbidimetric assay and normalised according to the amount of protein detected in Western Blot. Representative results from one of three experiments are shown. Bars represent the mean±SD of triplicates. **p-value<0.01 by unpaired two-tailed T-test.

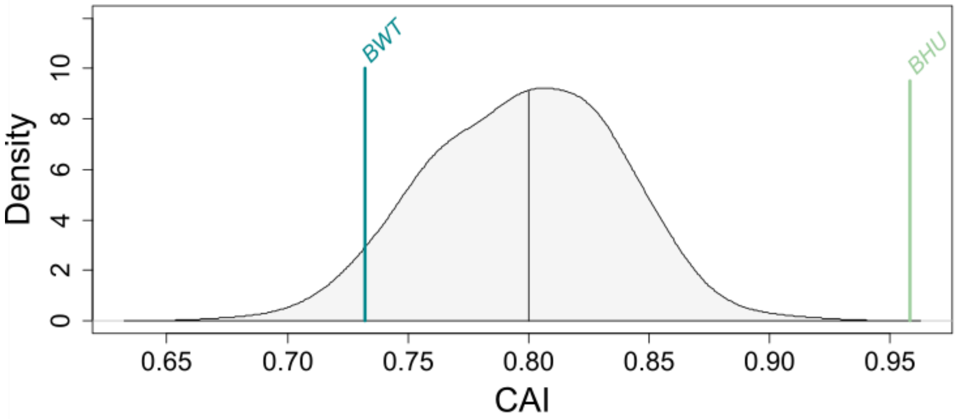


**Supplementary Fig. 5:**

CAI analysis of the wild-type hyaluronidase (*BWT*) and the codon humanised bee hyaluronidase (*BHU*), in relation to human genes CAI.

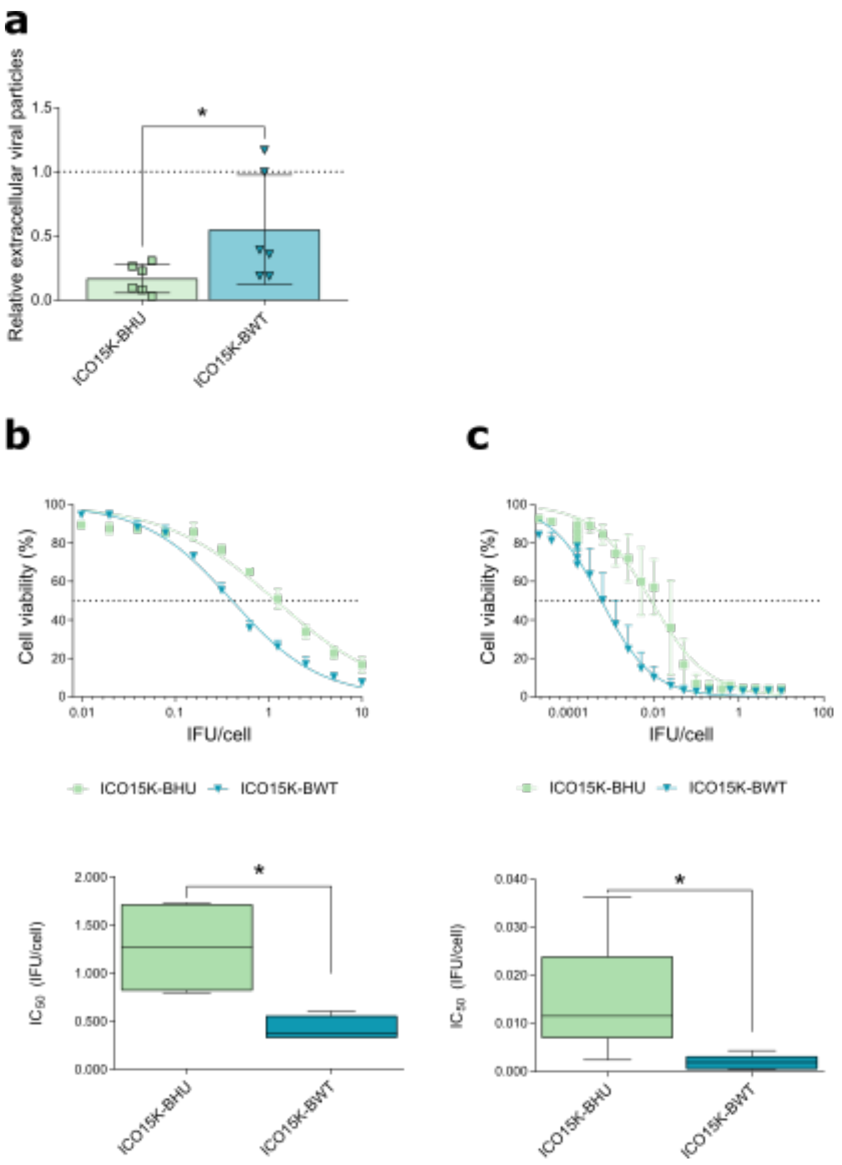


**Supplementary Fig. 6: Codon humanised bee hyaluronidase transgene (*BHU*) impairs viral oncolytic activity.**

**a** qPCR relative quantitation of the extracellular viral particles released to the supernantants of MIA PaCa-2 cells infected with 5 IFU of VCN-01, ICO15K-BHU or ICO15K-BWT at 72 hpi,. The dashed line represents VCN-01 values. Data is represented as the mean ±SEM; each dot corresponds to an independent experimental replicate. *p<0.05 (two tailed Mann- Whitney test).

**b-c** *In vitro* oncolytic activity assay in MIA PaCa-2 **(b)**and NP18 *(c)* cells. Cells were infected with a dose range of ICO15K-BHU and ICO15K-BWT and their viability was measured 7 days PI by MTT assay. Viability curves are represented in the upper panel. Data is represented as mean ±SEM for at least four independent experiments. The lower panel shows $IC_{50}$ values calculated from viability curves. Data is represented as box plot of four independent experiments. *p<0.05 (two tailed Mann- Whitney test).

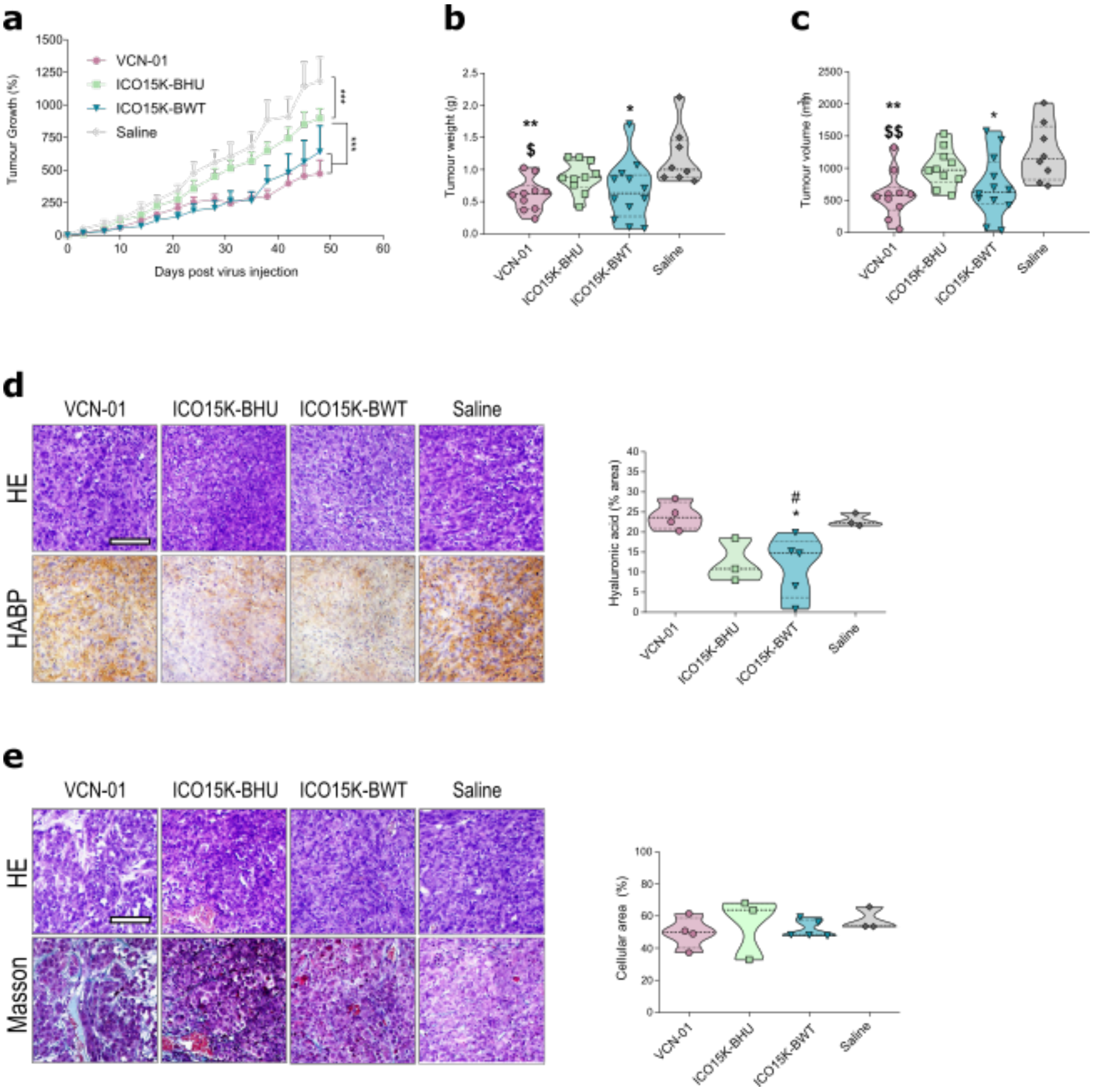


**Supplementary Fig. 7: Modulating codon optimization of bee hyaluronidase transgene leads to optimal therapeutic index response in MIA PaCa-2 *in vivo* model.**

**a** *In vivo* tumour growth assay in mice bearing subcutaneous MIA PaCa-2 tumours. Animals were intravenously treated with saline solution or $4{\cdot}10^{10}$vp/animal of either VCN-01, ICO15K-BHU, ICO15K-BWT or ICO15K-BAd (n≥6 animals/group). Follow-up of tumour volumes is represented as mean of percentage of growth ±SEM.

**b** Tumour weight at end-point.

**c** Tumour volume at end-point.

**d** HA acid staining quantitation of deparaffinised tumour sections with HABP. Scale bar 100 µm. Left panels: representative images of HA staining. Right panels: HA acid quantification representing stained area in at least 5 fields from the tumour sections of at least 3 different mice.

**e** Tumour cellularity measured by Masson staining. Scale bar 100 µm. Left panels: representative images of HA staining. Right panels: HA acid quantification.

*p<0.05, **p<0.01, ***p<0.001, # p<0.05, ## p<0.01, $ p<0.05, $$ p<0.01 * represents statistical differences in relation to saline group; # represents statistical differences in relation to saline VCN-01 group, $ represents statistical differences in relation to ICO15K-BHU group.

Fig. 1g

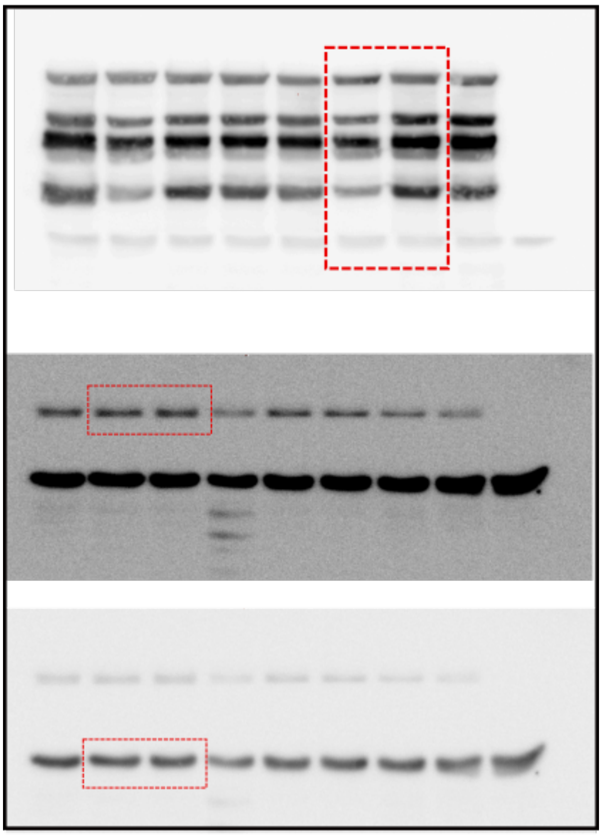

Fig. 2e

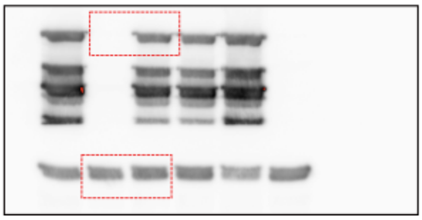

**Supplementary Fig. 8: Uncropped images of western blot figures shown in the main paper.**

**Supplementary Table 1. Primer sequences**

| Primer set | Primer name | Primer Sequence |
|---|---|---|
| 1 | Fw_BamHI_EGFP | TATGCGGATCCATGGTGAGCAAGGGCGAGGAGC |
| | Rv_EcoRI_EGFP | TATGCGAATTCTTACTTGTACAGCTCGTCCATGC |
| 2 | Fw_BamHI_pMONO-neo-GFP | TATGCGGATCCATGAGCAAGGGAGAAGAACTC |
| | Rv_EcoRI_pMONO-neo-GFP | TATGCGAATTCTTACTTGTACAGCTCATCCATTCC |
| 3 | sPH20_Fwd | CCACCGGTGCCACCATGGG |
| | sPH20_Rev | AACGCGGCCGCTTTATTAGTGGTGGTGGTGGTGGTGGGGGCCCTGGAACAGCACCTCCAGAGATAGTGTGGAGGGTGAAGC |
| 4 | BHyal_Fwd | ATCCACCGGTCCACCATGTCTCGGCCTCTCGTGAT |
| | BHyal_Rv | CGCGGCGGCCGCTTTAGTGGTGGTGGTGGTGGTGGGGGCCCTGGAACAGCACCTCCAGCACTTGGTCCACGCTCACGTC |

| | | |
|---|---|---|
| **5** | Fw_Ad_late_pMONO-neo-GFP | CGTGTTTATTTTTCAATTGGTACTAAGCGGTGATGTTTCTGATCAGCCACCATGAGCAAGGGAGAAGAACTC |
| | Rv_Ad_late_pMONO-neo-GFP | GAATGAAAAATGACTTGAAATTTTCTGCAATTGAAAAATAAAGTTTATTATTACTTGTAC AGCTCATCCATTCC |
| **6** | Recombination BHU FW | ATCGTTTGTGTTATGTTTCAACGTGTTTATTTTTCAATTGGTACTAAGCGGTGATGTTTCTGATCAGCCACCATGTCTCGGCCTCTCGTGA |
| | Recombination BHU RV | GCTATACTACTGAATGAAAAATGACTTGAAATTTTCTGCAATTGAAAAATAAAGTTTATTACACTTGGTCCACGCTCA |
| **7** | Fiber-GFP Fw | CAATTGGTACTAAGCGGTGATGTTTCTGATCAGCCACCATGGTGAGCAAGGGCGAGG |
| | Fiber-GFP Rv | gacttgaaattttctgcaattgaaaaataaagtttattaCTTGTACAGCTCGTCCATGC |

| | | |
|---|---|---|
| **8** | E4 miARN cua Fw | GAAAACTACAATTCCCAACACATACAAGTTACTCCG CCCTAACTTTTATTTTATCGAATCT GC |
| | E4 miARN cua Rv | CGTGGCGCGGGGCGTGGGAACGGGGCGGGTGAC GTAGGTTACATTGATTATTGACTAG |
| **9** | qPCR-Ad-genome-Fw | GCCGCAGTGGTCTTACATGCACATC |
| | qPCR-Ad-genome-Rv | CAGCACGCCGCGGATGTCAAAG |
| **10** | qPCR-hexon-Fw | GTCTACTTCGTCTTCGTTGTC |
| | qPCR-hexon-Rv | TGGCTTCCACGTACTTTG |
| **11** | qPCR-fiber-Fw | CTCCAACTGTGCCTTTTC |
| | qPCR-fiber-Rv | GGCTCACAGTGGTTACATT |
| **12** | qPCR-ACTB-Hs-Fw | CTGGAACGGTGAAGGTGACA |
| | qPCR-ACTB-Hs-Rv | GGGAGAGGACTGGGCCATT |

### Supplementary Data 1. GFP sequence alignments

#### Gene sequence alignment

```
EGFP            ATGGTGAGCAAGGGCGAGGAGCTGTTCACCGGGGTGGTGCCCATCCTGGTCGAGCTGGAC    60
LGFP            ---ATGAGCAAGGGAGAAGAACTCTTTACTGGTGTTGTCCCAATTCTGGTTGAGCTGGAT    57
                    ********** ** ** ** ** ** ** ** ** ** ** ***** ********

EGFP            GGCGACGTAAACGGCCACAAGTTCAGCGTGTCCGGCGAGGGCGAGGGCGATGCCACCTAC    120
LGFP            GGTGATGTGAATGGCCACAAATTCTCTGTGTCTGGTGAAGGTGAAGGAGATGCAACTTAT    117
                ** ** ** ** ******** ***   ***** ** ** ** ** ** ***** ** **

EGFP            GGCAAGCTGACCCTGAAGTTCATCTGCACCACCGGCAAGCTGCCCGTGCCCTGGCCCACC    180
LGFP            GGAAAGCTGACTCTGAAGTTCATTTGTACAACAGGAAAGCTGCCAGTGCCTTGGCCAACT    177
                ** ******** *********** ** ** ** ** ******** ***** ***** **

EGFP            CTCGTGACCACCCTGACCTACGGCGTGCAGTGCTTCAGCCGCTACCCCGACCACATGAAG    240
LGFP            CTGGTGACCACCCTGACTTATGGTGTTCAATGTTTCAGCAGGTACCCTGACCACATGAAG    237
                ** ************** ** ** ** ** ** ****** * ***** ************

EGFP            CAGCACGACTTCTTCAAGTCCGCCATGCCCGAAGGCTACGTCCAGGAGCGCACCATCTTC    300
LGFP            CAGCATGACTTCTTTAAATCTGCAATGCCAGAAGGTTATGTTCAGGAGAGGACAATCTTC    297
                ***** ******** ** ** ** ***** ***** ** ** ****** * ** ******

EGFP            TTCAAGGACGACGGCAACTACAAGACCCGCGCCGAGGTGAAGTTCGAGGGCGACACCCTG    360
LGFP            TTTAAGGATGATGGAAATTATAAGACAAGGGCAGAAGTGAAGTTTGAAGGTGATACACTG    357
                ** ***** ** ** ** ** *****  * ** ** ******** ** ** ** ** ***

EGFP            GTGAACCGCATCGAGCTGAAGGGCATCGACTTCAAGGAGGACGGCAACATCCTGGGGCAC    420
LGFP            GTTAACAGAATTGAGCTGAAAGGCATTGATTTTAAGGAAGATGGAAACATTCTGGGTCAC    417
                ** *** * ** ******** ***** ** ** ***** ** ** ***** ***** ***

EGFP            AAGCTGGAGTACAACTACAACAGCCACAACGTCTATATCATGGCCGACAAGCAGAAGAAC    480
LGFP            AAGCTGGAGTACAACTATAATTCTCACAATGTTTACATTATGGCAGATAAGCAGAAGAAT    477
                ***************** **    ***** ** ** ** ***** ** ***********

EGFP            GGCATCAAGGTGAACTTCAAGATCCGCCACAACATCGAGGACGGCAGCGTGCAGCTCGCC    540
LGFP            GGAATTAAGGTTAATTTCAAGATTAGACACAACATTGAGGATGGATCTGTCCAACTGGCA    537
                ** ** ***** ** ********  * ******** ***** **    ** ** ** **

EGFP            GACCACTACCAGCAGAACACCCCCATCGGCGACGGCCCCGTGCTGCTGCCCGACAACCAC    600
LGFP            GACCATTACCAGCAGAACACCCCTATTGGTGATGGCCCAGTTCTCCTCCCAGATAATCAC    597
                ***** **************** ** ** ** ***** ** ** ** ** ** ** ***

EGFP            TACCTGAGCACCCAGTCCGCCCTGAGCAAAGACCCCAACGAGAAGCGCGATCACATGGTC    660
LGFP            TATCTCCGCACTCAATCTGCTCTGTCCAAAGACCCTAATGAGAAAAGAGACCACATGGTC    657
                ** **  **** ** ** ** ***  ********* ** *****  * ** *********

EGFP            CTGCTGGAGTTCGTGACCGCCGCCGGGATCACTCTCGGCATGGACGAGCTGTACAAGTAA    720
LGFP            CTCCTGGAGTTTGTGACAGCAGCAGGAATTACTCTGGGAATGGATGAGCTGTACAAGTAA    717
                ** ******** ***** ** ** ** ** ***** ** ***** **************
```

#### Protein sequence alignment

```
EGFP            MVSKGEELFTGVVPILVELDGDVNGHKFSVSGEGEGDATYGKLTLKFICTTGKLPVPWPT 60
LGFP            -MSKGEELFTGVVPILVELDGDVNGHKFSVSGEGEGDATYGKLTLKFICTTGKLPVPWPT 59
                 :**********************************************************

EGFP            LVTTLTYGVQCFSRYPDHMKQHDFFKSAMPEGYVQERTIFFKDDGNYKTRAEVKFEGDTL 120
LGFP            LVTTLTYGVQCFSRYPDHMKQHDFFKSAMPEGYVQERTIFFKDDGNYKTRAEVKFEGDTL 119
                ************************************************************

EGFP            VNRIELKGIDFKEDGNILGHKLEYNYNSHNVYIMADKQKNGIKVNFKIRHNIEDGSVQLA 180
LGFP            VNRIELKGIDFKEDGNILGHKLEYNYNSHNVYIMADKQKNGIKVNFKIRHNIEDGSVQLA 179
                ************************************************************

EGFP            DHYQQNTPIGDGPVLLPDNHYLSTQSALSKDPNEKRDHMVLLEFVTAAGITLGMDELYK* 239
LGFP            DHYQQNTPIGDGPVLLPDNHYLRTQSALSKDPNEKRDHMVLLEFVTAAGITLGMDELYK* 238
                ********************** *************************************
```

### Supplementary Data 2. Hyaluronidase sequence alignments

### Gene sequence alignment

```
BHU   ATGAGCAGACCCCTGGTGATCACCGAGGGCATGATGATCGGCGTGCTGCTGATGCTGGCC 60
BWT   ATGTCTCGGCCTCTCGTGATCACGGAAGGGATGATGATTGGAGTGTTGCTAATGCTAGCC 60
      ***     * ** ** ******** ** ** ******** ** *** **** ***** ***

BHU   CCTATCAACGCCCTGCTGCTGGGCTTTGTGCAGAGCACTCCCGACAACAACAAGACCGTG 120
BWT   CCGATAAACGCGTTATTACTCGGCTTCGTACAGAGCACCCCCGACAACAACAAAACCGTA 120
      ** ** *****  *  * ** ***** ** ******** ************** *****

BHU   CGCGAGTTTAACGTGTACTGGAACGTGCCCACCTTCATGTGCCACAAGTACGGCCTGCGC 180
BWT   CGGGAGTTCAACGTTTACTGGAACGTGCCCACCTTTATGTGCCATAAATACGGGCTACGG 180
       ** ***** ***** ******************* ******** ** ***** ** **

BHU   TTTGAGGAGGTGAGCGAGAAGTACGGCATCCTGCAGAACTGGATGGACAAGTTTCGCGGA 240
BWT   TTCGAAGAAGTATCGGAGAAATATGGTATTCTACAGAACTGGATGGATAAGTTTCGGGGC 240
      ** ** ** **    ***** ** ** ** ** ************** ******** **

BHU   GAGGAGATTGCCATCCTGTACGACCCTGGCATGTTTCCCGCTCTGCTGAAGGATCCCAAC 300
BWT   GAGGAGATCGCGATCCTTTACGACCCTGGAATGTTCCCGGCGTTGCTGAAAGACCCGAAT 300
      ******** ** ***** *********** ***** ** **  ******* ** ** **

BHU   GGCAACGTGGTGGCTCGCAACGGAGGCGTGCCTCAGCTGGGCAACCTGACCAAGCACCTG 360
BWT   GGGAACGTGGTGGCGAGGAACGGCGGTGTCCCGCAACTGGGCAATCTCACCAAGCATCTG 360
      ** ***********  * ***** ** ** ** ** ******** ** ******** ***

BHU   CAGGTGTTTCGCGACCACCTGATCAACCAGATTCCCGACAAGAGCTTTCCCGGAGTGGGC 420
BWT   CAAGTATTTCGGGACCACTTGATCAATCAGATCCCGGACAAGTCGTTTCCCGGCGTGGGG 420
      ** ** ***** ****** ******* ***** ** ******    ******** *****

BHU   GTGATCGACTTTGAGAGCTGGAGACCCATCTTTCGCCAGAACTGGGCTAGCCTGCAGCCC 480
BWT   GTGATCGATTTCGAAAGTTGGAGGCCGATATTCAGACAGAACTGGGCCTCCCTCCAGCCT 480
      ******** ** ** ** ***** ** ** **  * ***********    *** *****

BHU   TACAAGAAGCTGAGCGTGGAGGTGGTGCGCAGAGAGCACCCTTTCTGGGACGACCAGCGC 540
BWT   TACAAGAAACTGTCCGTAGAGGTGGTTCGCCGTGAGCATCCGTTCTGGGACGATCAGAGG 540
      ******** ***  *** ******** *** * ***** ** *********** *** *

BHU   GTGGAGCAGGAGGCCAAGAGACGCTTTGAGAAGTACGGCCAGCTGTTCATGGAGGAGACC 600
BWT   GTGGAGCAGGAGGCGAAACGAAGGTTCGAGAAATACGGGCAGCTTTTCATGGAGGAGACG 600
      ************** **  ** * ** ***** ***** ***** **************

BHU   CTGAAGGCAGCCAAGCGCATGAGACCTGCTGCCAACTGGGGCTACTACGCCTACCCTTAC 660
BWT   TTGAAAGCGGCGAAACGGATGAGGCCGGCCGCCAATTGGGGATACTACGCCTACCCTTAT 660
       **** ** ** ** ** ***** ** ** ***** ***** ****************

BHU   TGCTACAACCTGACTCCCAACCAGCCCAGCGCCCAGTGCGAGGCCACTACCATGCAGGAG 720
BWT   TGCTACAATCTGACGCCGAATCAGCCGAGCGCCCAATGCGAAGCGACCACCATGCAGGAG 720
     ******** ***** ** ** ***** ******** ***** ** ** ************

BHU   AACGACAAGATGAGCTGGCTGTTTGAGAGCGAGGACGTGCTGCTGCCCAGCGTGTACCTG 780
BWT   AACGATAAAATGTCGTGGCTGTTCGAGTCGGAAGACGTCCTCCTTCCGTCCGTTTACTTG 780
      ***** ** ***   ******** ***    ** ***** ** ** **    *** *** **

BHU   CGCTGGAACCTGACCAGCGGCGAGCGCGTTGGACTGGTTGGAGGACGCGTGAAGGAGGCC 840
BWT   AGATGGAATCTGACGAGCGGCGAAAGAGTGGGCCTGGTCGGTGGCCGCGTGAAGGAGGCG 840
       * ***** ***** ********  * ** ** ***** ** ** **************

BHU   CTGAGAATTGCCAGACAGATGACCACTAGCCGCAAGAAGGTGCTGCCCTACTACTGGTAC 900
BWT   TTGAGAATAGCGAGGCAAATGACGACCAGCAGGAAGAAGGTTCTACCATATTACTGGTAC 900
       ******* ** ** ** ***** ** *** * ******** ** ** ** *********

BHU   AAGTACCAGGACAGACGCGACACCGACCTGAGCAGAGCCGACCTGGAAGCCACTCTGCGC 960
BWT   AAATATCAGGATCGAAGGGACACGGATTTGAGCAGGGCTGACCTCGAGGCAACTTTACGA 960
      ** ** *****  ** * ***** **  ******* ** ***** ** ** *** * **

BHU   AAGATCACCGACCTGGGAGCTGACGGCTTCATCATCTGGGGCAGCAGCGACGACATCAAC 1020
BWT   AAAATCACGGACCTCGGCGCCGACGGGTTCATCATTTGGGGAAGTTCCGACGATATAAAC 1020
      ** ***** ***** ** ** ***** ******** ***** **    ****** ** ***

BHU   ACCAAGGCCAAGTGCCTGCAGTTTCGCGAGTACCTGAACAACGAGCTTGGACCTGCCGTG 1080
```

```
BWT   ACGAAGGCGAAGTGCCTACAATTCAGGGAATACCTGAACAACGAGTTGGGCCCTGCCGTT 1080
      ** ***** ******** ** **  * ** *************** * ** ********

BHU   AAGCGCATTGCCCTGAACAACAACGCCAACGACAGACTGACCGTGGACGTGAGCGTGGAC 1140
BWT   AAACGAATCGCGTTGAACAACAACGCGAACGATCGACTGACGGTGGACGTGAGCGTGGAC 1140
      ** ** ** **  ************* *****  ******* ******************

BHU   CAGGTGTAA 1149
BWT   CAAGTGTGA 1149
      ** **** *
```

### Protein sequence alignment

```
BHU   MSRPLVITEGMMIGVLLMLAPINALLLGFVQSTPDNNKTVREFNVYWNVPTFMCHKYGLR 60
BWT   MSRPLVITEGMMIGVLLMLAPINALLLGFVQSTPDNNKTVREFNVYWNVPTFMCHKYGLR 60
      ************************************************************
BHU   FEEVSEKYGILQNWMDKFRGEEIAILYDPGMFPALLKDPNGNVVARNGGVPQLGNLTKHL 120
BWT   FEEVSEKYGILQNWMDKFRGEEIAILYDPGMFPALLKDPNGNVVARNGGVPQLGNLTKHL 120
      ************************************************************

BHU   QVFRDHLINQIPDKSFPGVGVIDFESWRPIFRQNWASLQPYKKLSVEVVRREHPFWDDQR 180
BWT   QVFRDHLINQIPDKSFPGVGVIDFESWRPIFRQNWASLQPYKKLSVEVVRREHPFWDDQR 180
      ************************************************************

BHU   VEQEAKRRFEKYGQLFMEETLKAAKRMRPAANWGYYAYPYCYNLTPNQPSAQCEATTMQE 240
BWT   VEQEAKRRFEKYGQLFMEETLKAAKRMRPAANWGYYAYPYCYNLTPNQPSAQCEATTMQE 240
      ************************************************************

BHU   NDKMSWLFESEDVLLPSVYLRWNLTSGERVGLVGGRVKEALRIARQMTTSRKKVLPYYWY 300
BWT   NDKMSWLFESEDVLLPSVYLRWNLTSGERVGLVGGRVKEALRIARQMTTSRKKVLPYYWY 300
      ************************************************************
```
